# Probing neuronal functions with precise and targeted laser ablation in the living cortex: reply

**DOI:** 10.1101/2022.04.15.488384

**Authors:** Zongyue Cheng, Jianian Lin, Yiyong Han, Bowen Wei, Baoling Lai, Baoming Li, Meng Cui, Wen-Biao Gan

## Abstract

In the comment on the article [Optica 8, 12 (2021)], the authors performed theoretical calculation to show that with a 1.1 NA objective and 140 fs laser pulses with single pulse energy of 2.2 nJ, and 80 MHz repetition rate, the focal point temperature rises 0.3 K and reaches equilibrium after 100 μs in water. They suggest that the damage to brain tissue by laser could not be caused by thermal effects but rather by plasma-mediated chemical effects. To quantify the thermal accumulation due to the fs laser illumination in living animals, we used a thermocouple sensor to measure the temperature change in the vicinity of the laser focus at a depth of 300 μm in the adult mouse cortex. Our results show that at 930 nm wavelength, 25 μm from the laser focus, 19-300 mW laser can all lead to a brain tissue temperature rise of more than 0.3 K at 1 second and the maximum equilibrium temperature rise of more than 28 K. These experimental measurements are significantly higher than theoretically calculated values in the comment. These results suggest that the thermal accumulation effect of focused low-energy pulses from fs laser oscillators could contribute significantly to the collateral damage in the living brain.

---

Our recently published article [1] focuses on the application of the amplified femtosecond (fs) laser system (AFL) to specifically ablate individual cells in the mammalian brain. We developed an iterative convergence power and Galvo control system, which were able to control precisely both the power of each amplified femtosecond laser pulse and the targeted ablation position within a 3D volume of the mouse cerebral cortex. The comment supports our conclusion that plasma-mediated targeted cell ablation by a single fs laser pulse is a versatile tool for studying the function of individual cells with minimal collateral damage in complicated neural circuits.

To highlight the accuracy of AFL, we used conventional Ti:sapphire fs laser oscillators for damage comparison in previous studies (Ref. [1] Fig. 3i-l, Supplement Fig. 4). We found that the conventional Ti:sapphire fs laser caused collateral damage to the surrounding tissue after various irradiation times. Specifically, in Fig. 3i-l (Ref. [1]), we used 2.2 nJ (average power of 176 mW) pulses for small ROI scans, in Supplement Fig. 4d,e (Ref. [1]), we used 3.75 nJ (average power of 300 mW) laser pulses for single spot irradiation. The results showed that the nucleus staining dye Hoechst 33342 decreased and propidium iodide increased in the targeted cells after irradiation, while the fluorescent intensity of YFP within surrounding structures decreased. In contrast to AFL ablation (Ref. [1] Fig. 3g,h, Supplement fig 4f), we suggest that this collateral damage to the adjacent tissues might be due to heat accumulation during the laser scanning. In the comment, the authors performed theoretical calculation which suggests that the focal point temperature increased by only 0.3 K at the pulse energy of 2.2 nJ (average laser power: 176 mW) and reached equilibrium within 100 μs. They concluded that tissue damage was not caused by this minor temperature increase, but rather by free-electron-mediated biomolecular damage related to ROS production.

To determine temperature change during conventional laser ablation under experimental conditions in Ref. [1], we directly measured the cortical temperature in living mouse cortex during laser irradiation. We inserted a 100 μm diameter thermocouple sensor (PerfectPrime, TL0201, 100 μm) into the primary sensory cortex of 2-month-old adult mice at a depth of approximately 300 μm, and took readings via thermometer (Fluke, 51-II). We parked the 80 MHz 930 nm laser beam with different intensities at a single spot for at least 6 seconds and measured the temperature of the living brain at various distances from the focal point. The experiments were performed using a Nikon 25X NA 1.1 water objective. To visualize the thermocouple sensor, we used Sulforhodamine 101 (Sigma-Aldrich) to stain the thermocouple sensor before insertion (Fig 1a-b).

**Fig.1.**
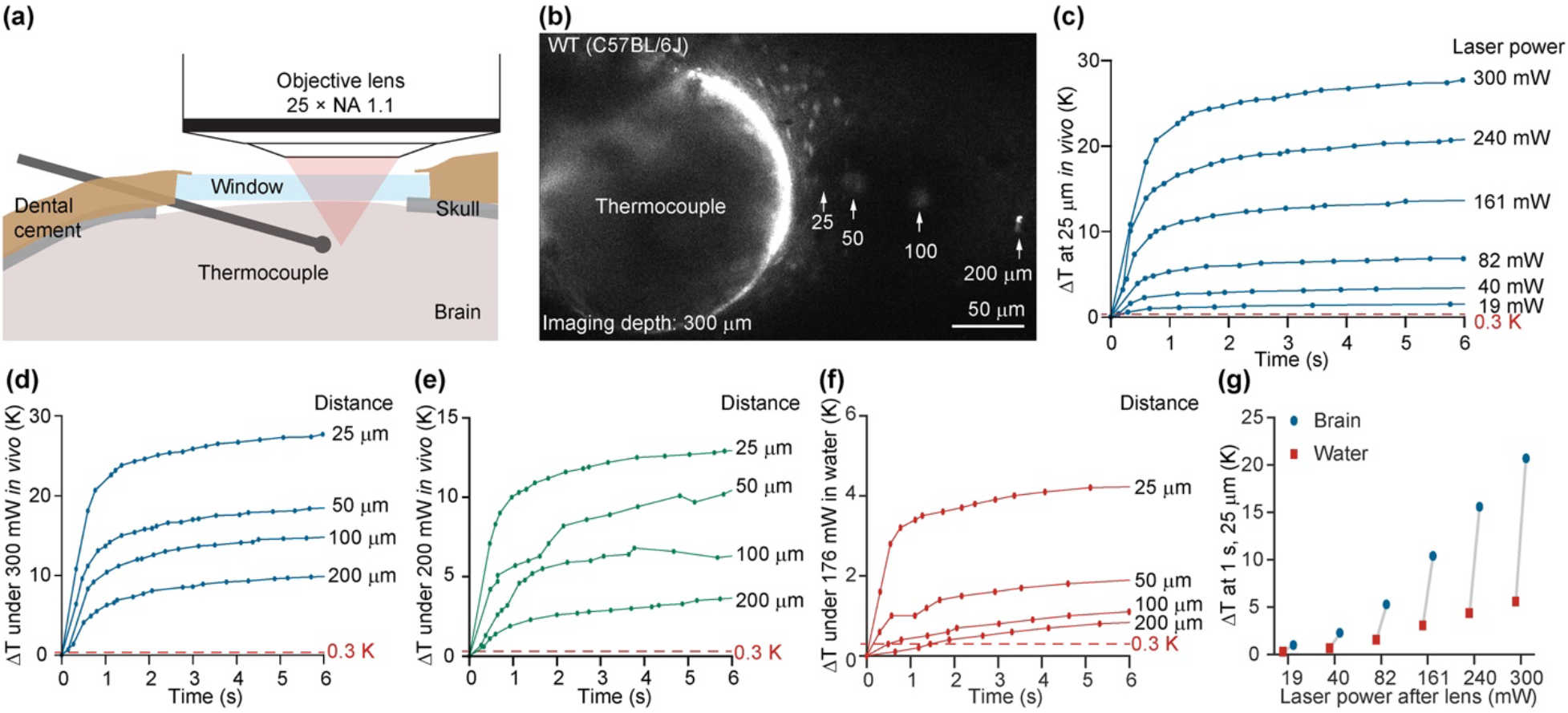
Thermocouple-based intracerebral temperature measurement in laser-irradiated mice. (a) Schematic diagram of thermocouple sensor measurement. (b) Two-photon imaging of the thermocouple sensor and laser irradiation positions. (c) Temperature changes in the living brain tissue caused by different light intensities at 25 μm from the laser focus. (d) Temperature changes in the living brain tissue induced by 300 mW laser at different distances. (e) Temperature changes in the live brain tissue induced by 200 mW laser at different distances. (f) Temperature changes in water induced by 176 mW laser at different distances. (g) Temperature changes at 1 s caused by different laser powers in the brain vs. water.

When the distance between the laser focus and the thermocouple sensor was kept at 25 μm, we found that the temperature rises at 1 s exceeded 0.3 K for all measured laser intensities, and the temperature equilibration time caused by different intensities exceeded 100 μs (Fig 1c). Furthermore, when we measured the temperature change from 25-200 μm from the focal point, we found that the laser with average power of 300 mW would cause a temperature change ranging from 10 to 28 K, and this increase was detectable at 200 μm (Fig 1d). At 200 mW, the temperature increased ~ 13 K at 25 μm and ~ 3 K at 200 μm (Fig 1e). We noticed that the authors of the comment used water as the medium for temperature diffusion calculation. We found that the temperature change in water according to the conditions simulated in the comment (80 MHz, 930 nm, 2.2 nJ, 140 fs laser pulse, water as diffusion medium) exceeded 0.3 K (Fig. 1f). In addition, when we compared the temperature change in water and live brain tissue, we found that the temperature change in water was significantly lower than in living brain tissue (Fig 1g) at various laser intensities (19-300 mW).

Based the findings above, we conclude that there is significant thermal accumulation around nearby tissues during focal irradiation with a conventional fs laser. The range of temperature elevation is consistent with the previous study [2–5]. Therefore, the thermal effects of low-energy pulses from a fs laser could contribute significantly to the collateral damage in the living brain. We do not exclude possible damaging effects caused by free-electron-mediated biomolecular damage due to ROS production as suggested by the authors in the comment. It would be interesting to test such a hypothesis in the future with the fluorescent ROS indicator in the living brain.

## Funding

ZC, JL, and MC were supported by NIH U01NS107689, U01NS118302, and R21EY032382.

## Acknowledgments

We thank all the members in the Cui Laboratory for their constructive discussion.

## Disclosures

The authors declare no conflict of interest.

## Data availability

Data underlying the results presented in this paper are not publicly available at this time but may be obtained from the authors upon reasonable request.

## Notes

### Competing Interest Statement

The authors have declared no competing interest.

